# Precursor of chemically expanded hepatocytes (pre-cHep) with 1-million-fold expansion potential and liver repopulation capacity

**DOI:** 10.64898/2026.05.15.725446

**Authors:** Linh My Huynh, Yuichiro Higuchi, Cherie Tsz-Yiu Law, Jakob Jeriha, Ines Battle, Rebecca Granskog, Shotaro Uehara, Fumihiko Kawamura, Victoria L Gadd, Tak Yung Man, Stuart J Forbes, Kosuke Yusa, Stephen Kwok-Wing Tsui, Hiroshi Suemizu, Keisuke Kaji

## Abstract

Primary human hepatocytes (PHHs) are the gold standard for toxicology and drug metabolism studies in industry. However, their limited availability, substantial batch-to-batch variability, and high cost restrict their use. Here, we report a novel culture condition that reprograms PHHs into a proliferative state. These proliferating cells, termed precursors of chemically expanded hepatocytes (pre-cHep), expand over 10^6^-fold within 30 days while retaining liver repopulation capacity comparable to PHHs. pre-cHep can further differentiate into chemically expanded hepatocytes (cHep) as three-dimensional (3D) spheroids within 7 days *in vitro*, exhibiting global gene expression profiles, albumin production, and cytochrome P450 (CYP) activities similar to 3D-cultured PHH spheroids (3D PHH). Efficient genetic manipulation of pre-cHep using CRISPR/Cas9 is also achievable. Together, pre-cHep and cHep represent a promising alternative to high-quality PHHs, providing a more affordable, reproducible, and scalable source of human hepatocytes for toxicology, drug metabolism studies, disease modelling, towards precision drug development.

## Introduction

Hepatocytes comprise approximately 80% of liver mass and play critical roles in numerous physiological processes, including detoxification, drug metabolism, regulation of lipid, glucose, and amino acid homeostasis, as well as the synthesis of coagulation factors and complement proteins. Fully functional human hepatocytes are therefore essential for a wide range of applications, including in vitro toxicology testing, drug screening, and disease modelling. Hepatocyte transplantation, facilitated by the ability to cryopreserve cells, has also been proposed as an alternative to whole-organ liver transplantation^1^, potentially overcoming key limitations such as limited and unpredictable organ availability and the risks associated with major surgery. However, the supply of primary human hepatocytes (PHHs) remains constrained, and their significant batch-to-batch variability and high cost hinder experimental optimization and standardization.

Although embryonic stem cells (ESCs) and induced pluripotent stem cells (iPSCs) have been explored as alternative sources, pluripotent stem cell-derived hepatocyte-like cells (PSC-HLCs) remain functionally immature and have yet to achieve a fully competent hepatocyte phenotype^2,3^. Similarly, induced hepatocytes (iHeps) generated through reprogramming of fibroblasts using hepatic transcription factors, as well as immortalized hepatocyte cell lines, exhibit limited hepatic functionality^4^.

Mature hepatocytes rarely proliferate in the healthy liver but can re-enter the cell cycle following injury, playing a central role in liver regeneration. However, when cultured in vitro, primary human hepatocytes (PHHs) rapidly lose most of their synthetic and metabolic functions within approximately one week under standard two-dimensional (2D) conditions. Previous studies have demonstrated that human hepatocytes can be reprogrammed into chemically induced liver progenitors (CLiPs) or CLiP-like cells (CLiPLCs) using small molecules^5–9^. Re-differentiated cells derived from CLiPs/CLiPLCs have shown promise as an alternative to PHHs^10^. However, maintaining their re-differentiation capacity during long-term in vitro expansion remains challenging. Notably, none of these cells have exhibited global gene expression profiles or metabolic activities comparable to primary hepatocytes after more than one month of expansion followed by differentiation^5–9^. As such, the large-scale production of functional human hepatocytes with consistent quality remains a major unmet goal.

More recently, the Sato laboratory reported the expansion of adult human hepatocytes as organoids, achieving approximately 10^6^-fold expansion over 100 days^11^. These human hepatocyte organoids (HHOs) displayed gene expression and metabolic functions closely resembling PHHs following 7-14 days of differentiation, even after extended in vitro expansion^11^. However, their in vivo liver repopulation capacity declined beyond approximately 3,000-fold expansion, even in the best-performing lines. In addition, their relatively slow proliferation rate (split ratio of 1:2-1:3 every two weeks) and reliance on Matrigel limit scalability and increase costs, posing challenges for industrial application.

Here, we report a novel 2D culture condition that reprograms PHHs into a proliferative state while preserving both liver repopulation capacity and in vitro re-differentiation potential for over 30 days, achieving up to 10^6^-fold expansion.

## Results

### Derivation of precursors of chemically expanded hepatocytes (pre-cHep) with L4 medium

Bipotent CLiPs/CLiPLCs reprogrammed from mature hepatocytes show strong potential to re-differentiate into hepatocyte-like cells^10^, but rapidly lose both their in vitro differentiation capacity and liver repopulation ability over time^5–9^. We hypothesized that this decline arises because these cells drift too far from the mature hepatocyte state during expansion. Maintaining a cellular state closer to mature hepatocytes during proliferation may therefore enable more efficient reversion to full functionality. Based on this rationale, we identified that forskolin and dibutyryl cyclic AMP (db-cAMP), previously used to promote maturation of hepatocyte-like cells^9,12,13^, enhance the expansion of human hepatocytes when combined with vactosertib, blebbistatin, CHIR9902, an HGF mimetic peptide (PG-001), and fetal bovine serum (FBS). CHIR9902, HGF, and FBS have been widely used in CLiP/CLiPLC culture systems^5–9^. Vactosertib is a potent oral ALK5/ALK4 inhibitor, functionally similar to the commonly used ALK4/5/7 inhibitor A83-01. Blebbistatin is a selective inhibitor of non-muscle myosin II ATPase that is a downstream effector of Rho-associated kinase (ROCK), and can substitute for the ROCK inhibitor Y-27632 in human pluripotent stem cell cultures^14^. Using this formulation (designated L2 medium), hepatocytes from young donors (e.g., J03, 10 months old; J06, 8 months old) exhibited sustained, near-linear proliferation for approximately 45 days, reaching up to 10^10^-fold expansion (Figure 1A). We termed these cells precursors of chemically expanded hepatocytes (pre-cHep). The importance of forskolin and db-cAMP for robust pre-cHep proliferation was confirmed by their removal (L1 medium) at day 20, which resulted in a marked reduction in proliferation rate (Figure 1A). Furthermore, screening of ten candidate compounds, identified from the literature^15–23^, we identified IL-6 faciliated clonal expansion of pre-cHep maintaining ALB expression when supplemented to L2 (L4 medium) (Figure 1B). In bulk populations, L4 enhanced pre-cHep proliferation compared to L2, with differences becoming pronounced after day 45 (Figures 1A and 1B). Cells cultured in L4 were smaller and displayed a more homogeneous morphology than those cultured in L2 and L1 (Figure 1C). In addition, L2- and L4-cultured pre-cHep retained the mature hepatocyte surface marker ASGR1 during proliferation, with L4 pre-cHep maintaining higher ASGR1 expression than L2 pre-cHep at later time points (Figure 1D). Removal of forskolin and db-cAMP from L2, yielding L1 medium, led to a more rapid loss of ASGR1 expression (Figure 1D). These data are consistent with the idea that forskolin, db-cAMP, and IL-6 help maintain pre-cHep in a state closer to mature hepatocytes. Importantly, when pre-cHep expanded for 60 days in L1, L2, or L4 were differentiated into chemically expanded hepatocytes (cHep) as 3D spheroids (Figure 1E), L4-derived cHep showed ALBUMIN production equivalent to that of 3D-cultured HepaSH, which are freshly isolated human hepatocytes expanded in the liver of TK-NOG-hIL6 mice^24^ (Figure 1F). L2-derived cHep and L1-derived cHep exhibited approximately 50% and less than 5% of the ALBUMIN production observed in HepaSH, respectively (Figure 1F). A similar trend was observed for CYP1A2, CYP2C9, and CYP3A4 activities, with L4- and L1-derived cHep showing the highest and lowest activities, respectively, although the absolute activities remained lower than those of 3D-cultured PHH spheroids (3D PHH) when using 60-day-expanded pre-cHep (Figure 1G). These findings indicate that forskolin, db-cAMP, and IL-6 are critical for preserving re-differentiation capacity in pre-cHep. Moreover, when ASGR1-positive L2/L4 pre-cHep were isolated at day 35 and expanded for an additional 25 days, the resulting cHep exhibited higher CYP activities than cHep derived from unenriched populations, reaching levels comparable to those of 3D PHH (Figure 1G). This suggests that ASGR1 is a useful marker for predicting re-differentiation capacity, and that isolating ASGR1-positive pre-cHep is an effective strategy for enriching cells with superior *in vitro* differentiation potential. A distinctive feature of previously reported CLiPs/CLiP-like cells is their ability to differentiate not only into hepatocytes but also into cholangiocytes under appropriate culture conditions^6,7^. In contrast, under the same conditions, pre-cHep expanded in L4 did not form epithelial structures with rhodamine 123 transport activity, an indicator of cholangiocyte function. They also failed to up-regulate cholangiocyte-associated genes, unlike human biliary epithelial cells (hBECs)^25^ (Extended Figure 1). Altogether, these data indicate that L4 pre-cHep are closer to mature hepatocytes and are distinct from previously reported CLiPs/CLiPLCs.

**Figure 1.**
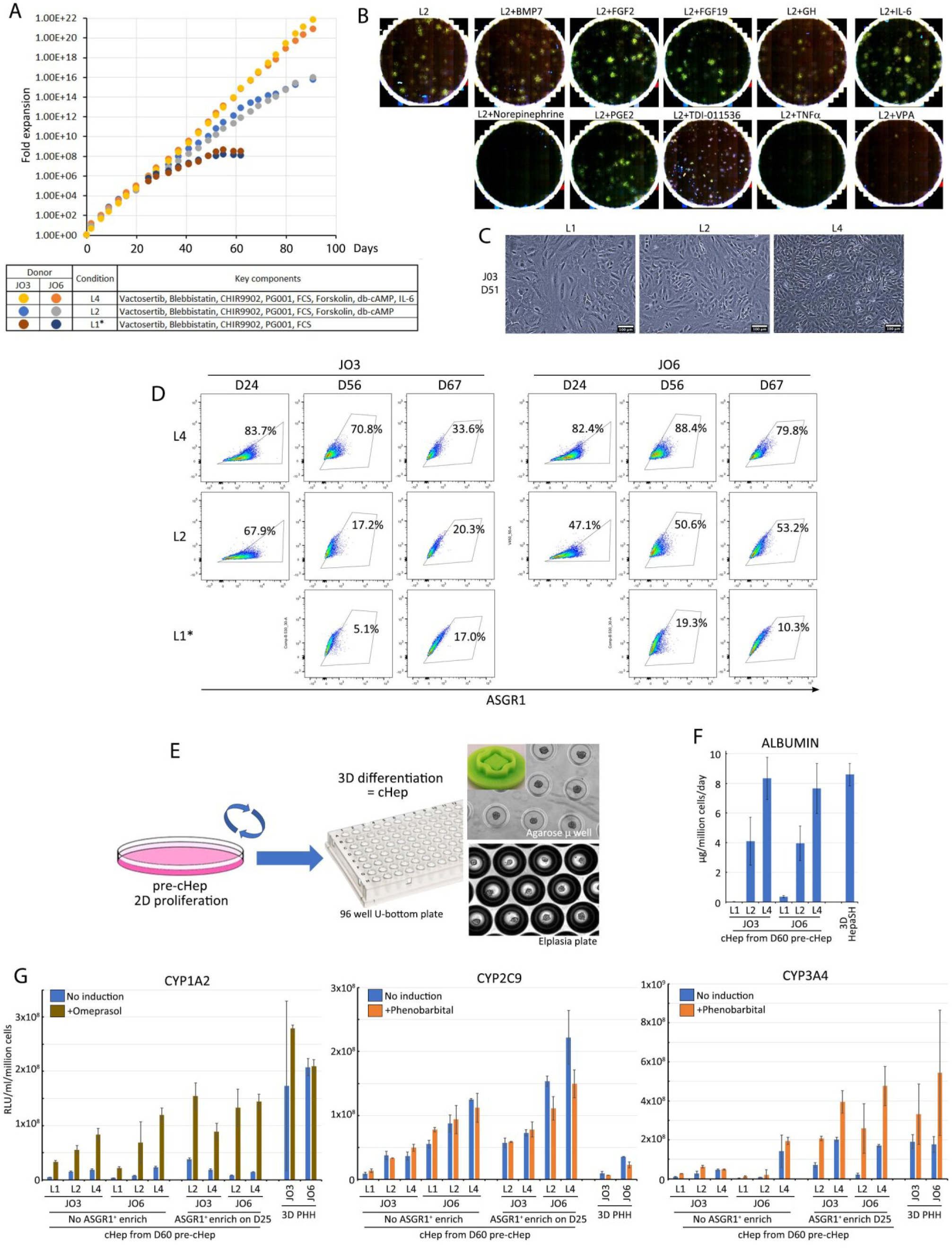
Expansion of pre-cHep lines with L4 medium. **A.** J03 and J06 PHHs were cultured in L2 or L4 medium, and cumulative cell numbers over passages were recorded. *Culture in L1 was introduced on day 20 by removing forskolin and db-cAMP from L2 and was stopped at day 76 due to minimal proliferation. Day 0 in this graph represents the first passage after 15 days of PHH culture. **B.** Mini-screen identifying IL-6 as an additional factor that improves L2 medium. L2 pre-cHep cells were seeded at clonal density in 6-well plates (500 cells/well) and cultured for 12 days under the indicated conditions before immunofluorescence staining for albumin. **C.** Representative images of pre-cHep cultured in L1, L2, and L4. **D.** Expression of ASGR1 in pre-cHep cultured in L1, L2, and L4 at the indicated time points during expansion. *Culture in L1 was introduced on day 20 by removing forskolin and db-cAMP from L2. **E.** Schematic diagram of 3D cHep generation from pre-cHep. **F.** Albumin secretion measured by ELISA in cHep generated from pre-cHep expanded for 60 days in L1, L2, or L4. **G.** CYP1A2, CYP2C9, and CYP3A4 activities in cHep generated from pre-cHep expanded for 60 days in L1, L2, or L4, with or without enrichment of ASGR1⁺ cells using PVLA at day 25. Data are presented as mean ± SD from technical duplicates.

### Liver repopulation capacity of L4 pre-cHep

Excitingly, three of the seven pre-cHep lines (J03, J06, and J13) established from different donor PHHs (Supplementary Table S1) in L4 medium retained excellent liver repopulation capacity even after approximately 30 days of expansion, corresponding to more than 10^6^-fold proliferation (∼20 doublings), following transplantation into TK-NOG-hIL6 mice (Figures 2A-2C). Among mice transplanted with day 30 J03 cells, 9 of 10 showed plasma cholinesterase (ChE) activity above 50 U/ml; similarly, 3 of 5 mice transplanted with day 30 J06 cells and 4 of 5 mice transplanted with day 28 J13 cells exceeded this threshold, which corresponds to approximately 50% liver repopulation in these animals (Figures 2A-2C). Immunohistochemistry further confirmed zone 3-biased expression of CYP3A4, as well as broader expression of CYP2C9 and, in repopulated livers, indicating functional integration of the transplanted human hepatocytes (Figure 2D). Healthy-looking hepatocytes distributed throughout the tissue, together with the absence of tumor formation after transplantation, are consistent with the lack of detectable copy number variation (CNV) changes between day 10 and day 60 pre-cHep by SNP array analysis (Extended Figure 2), indicating genetic stability during pre-cHep expansion. Interestingly, all pre-cHep lines that showed robust liver repopulation capacity—namely day 30 L4 J03, day 30 L4 J06, and day 28 L4 J13—contained a high proportion of ASGR1-positive cells (>50%) (Figures 1D and 2E). By contrast, after 60 days of culture (approximately 10^12^-fold expansion), both the J03 and J06 lines showed markedly reduced liver repopulation capacity, with 0 of 6 and only 1 of 5 transplanted mice, respectively, achieving cholinesterase activity above 50 U/ml. This loss of repopulation capacity correlated with a reduction in ASGR1 expression level (Figure 1D). Together, these findings suggest that strong ASGR1 expression is a useful indicator of pre-cHep liver repopulation capacity.

**Figure 2.**
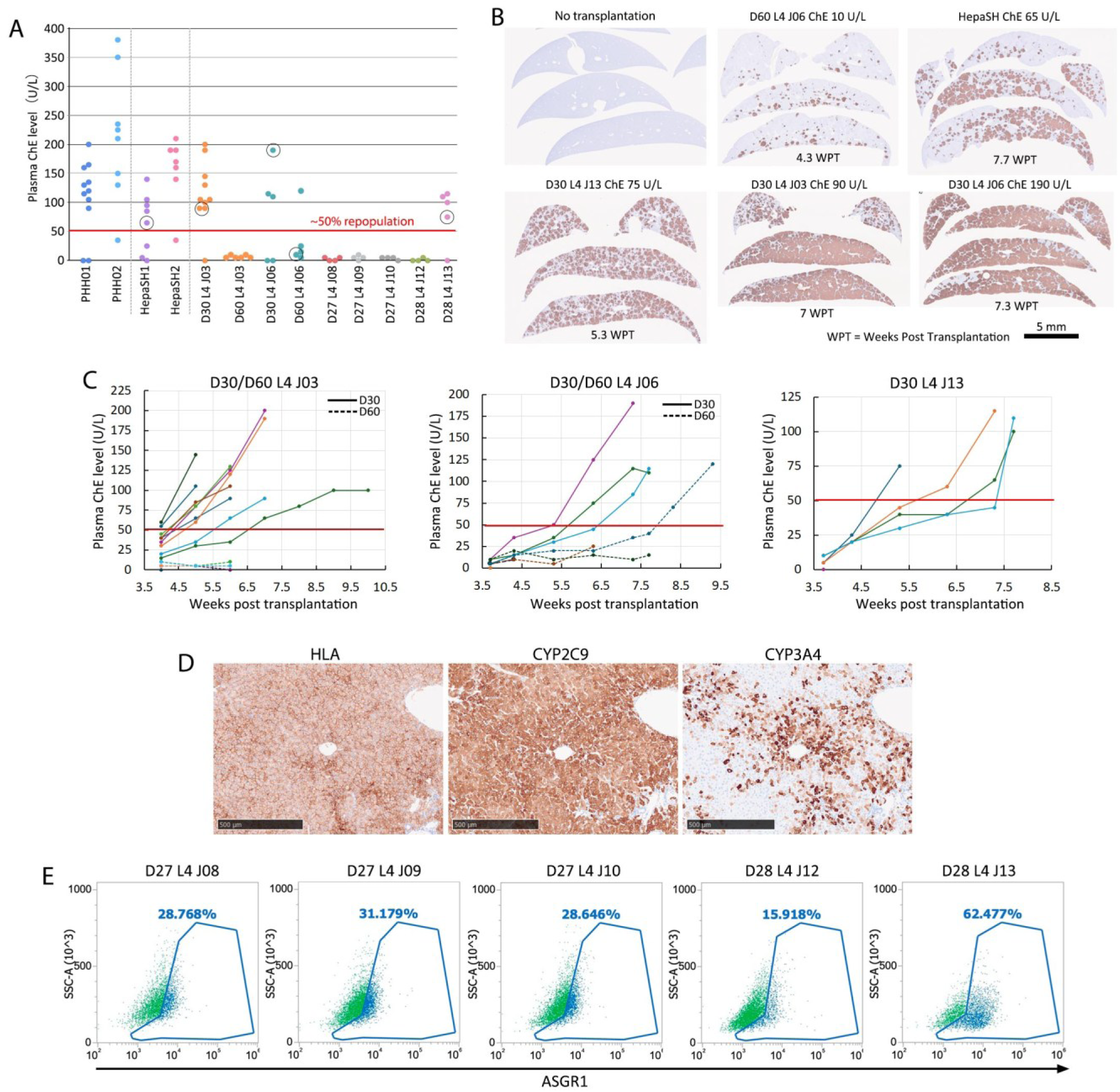
pre-cHep liver repopulation. **A.** J03, J06, J08, J09, J10, J12, and J13 pre-cHep lines expanded ∼10⁶-fold (30 days of culture) or ∼10¹²-fold (60 days of culture) in L4 medium were transplanted into TK-NOG hIL-6 mice. Plasma cholinesterase (ChE) activity measured at the final sampling time point for each mouse is shown. PHH and HepaSH transplantation served as positive controls. A ChE level of 50 U/mL corresponds to approximately 50% liver repopulation. **B.** Immunostaining for human mitochondria in liver sections from mice indicated by circles in panel A and control with no transplantation. **C.** Time course of plasma ChE levels up to the final sampling point or time of death in mice transplanted with J03, J06, and J13 pre-cHep. **D.** Immunostaining for HLA, CYP2C9, and CYP3A4 on serial liver sections following repopulation with J03 cells. **E.** ASGR1 expression in J08, J09, J10, J12, and J13 pre-cHep at approximately day 30 of expansion in L4. Data represent one of two independent experiments.

### cHep as a scalable alternative to 3D spheroid PHH

Next, we examined the global gene expression profiles of J03, J06, and J13 cHep differentiated from 30-day-cultured pre-cHep as spheroids, hereafter referred to as D30 cHep. The optimized differentiation protocol involved direct seeding of cryopreserved pre-cHep into Elplasia microwell plates at a density of 100 cells per microwell or into 96-well U bottom plates at a density of 1000 cells per well (Figure 3A). This is followed by 3 days of differentiation in re-differentiation medium R1 (L2 supplemented with IL-6, MEK inhibitor PD0325901, EGFR inhibitor Erlotinib, γ-secretase inhibitor DAPT, PI3K inhibitor LY294002), and then 4 days of maturation in serum-free maturation/maintenance BVCP medium supplemented with <u>v</u>actosertib, <u>b</u>lebbistatin, <u>C</u>HIR9902, and <u>P</u>D0325901 (Figure 3A). The use of these small molecules was informed by prior literature^26–29^ and iterative optimization. This 1-week culture period was comparable to the time required for cryopreserved PHHs to form spheroids (3D PHH) using a commercial medium (Figure 3A). Unlike PHHs, pre-cHep showed minimal cell death after thawing and formed more uniform spheroids (Figure 3B). When the RNA-seq data were integrated with published datasets^9,10,30,31^, the similarity between D30 cHep and 3D PHH became evident (Figure 3C) (Normalized log2 read counts in Supplementary Table S2). Other cell types that clustered with 3D PHH in principal component analysis (PCA) included previously reported hepatocyte-derived hepatocyte-like cells (Hep-HLCs), such as re-differentiated CLiPLCs reported by Zhang (ProliHH, passage 2 but not passage 6)^9^, Fu (HepLPC, passage 5)^5^, Kim (hCdH, passage number not indicated)^7^, and PHHs cultured in 2D with 5C medium^29^ (Figure 3C). This is consistent with previous reports that ProliHH, HepLPC, and hCdH retained liver repopulation capacity at these early passages, typically less than 10,000-fold expansion from PHHs. 5C medium maintains hepatocyte function in 2D without supporting proliferation, and the day 27 (D27) sample was more distant from PHH than the day 15 (D15) sample (Figure 3C). Notably, 3D-differentiated HepaRG cells, which are currently among the best PHH alternatives for toxicology testing, were even farther from 3D PHH in the PCA plot (Figure 3C). Hierarchical clustering and gene expression heatmap analysis of 264 liver-enriched genes from Human Protein Atlas (gene list in Supplementary Table S3) also showed that D30 cHep clustered closely with 3D PHH, although expression of some genes remained lower than in uncultured PHH and liver samples (Figure 3D). Distance-based similarity scoring of this gene set likewise identified D30 cHep as exhibiting the highest similarity to Liver/PHH among all cultured samples (Extended Figure 3A). Distance-based similarity scoring, hierarchical clustering, and gene-expression heatmaps of KEGG and GO gene sets linked to key liver functions, including xenobiotic metabolism, amino-acid biosynthesis, bile-acid biosynthesis, the urea cycle, fatty-acid metabolism, retinol metabolism, and the complement and coagulation cascade (gene lists in Supplementary Table S3), showed that D30 cHep and 3D PHH cluster closely together and are consistently among the samples most similar to liver/PHH (Figures 3E-3H, Extended Figures 3 and 4). In addition to RNA expression profiling, immunofluorescence and cholyl-lysyl-fluorescein (CLF) staining confirmed robust expression of ALBUMIN, HNF4A, CYP2E1, ASGR1, and bile canaliculi formation in cHep (Figures 3I and 3J). Given the rapid, large-scale expansion of pre-cHep in simple 2D culture without Matrigel, their straightforward and rapid differentiation from cryopreserved pre-cHep into mature cHep in a format comparable to standard 3D PHH culture, and their global gene-expression profile resembling that of 3D PHH, pre-cHep/cHep are well suited for applications that currently rely on 3D PHH and for contexts where PHH cost and variability have previously limited use.

**Figure 3.**
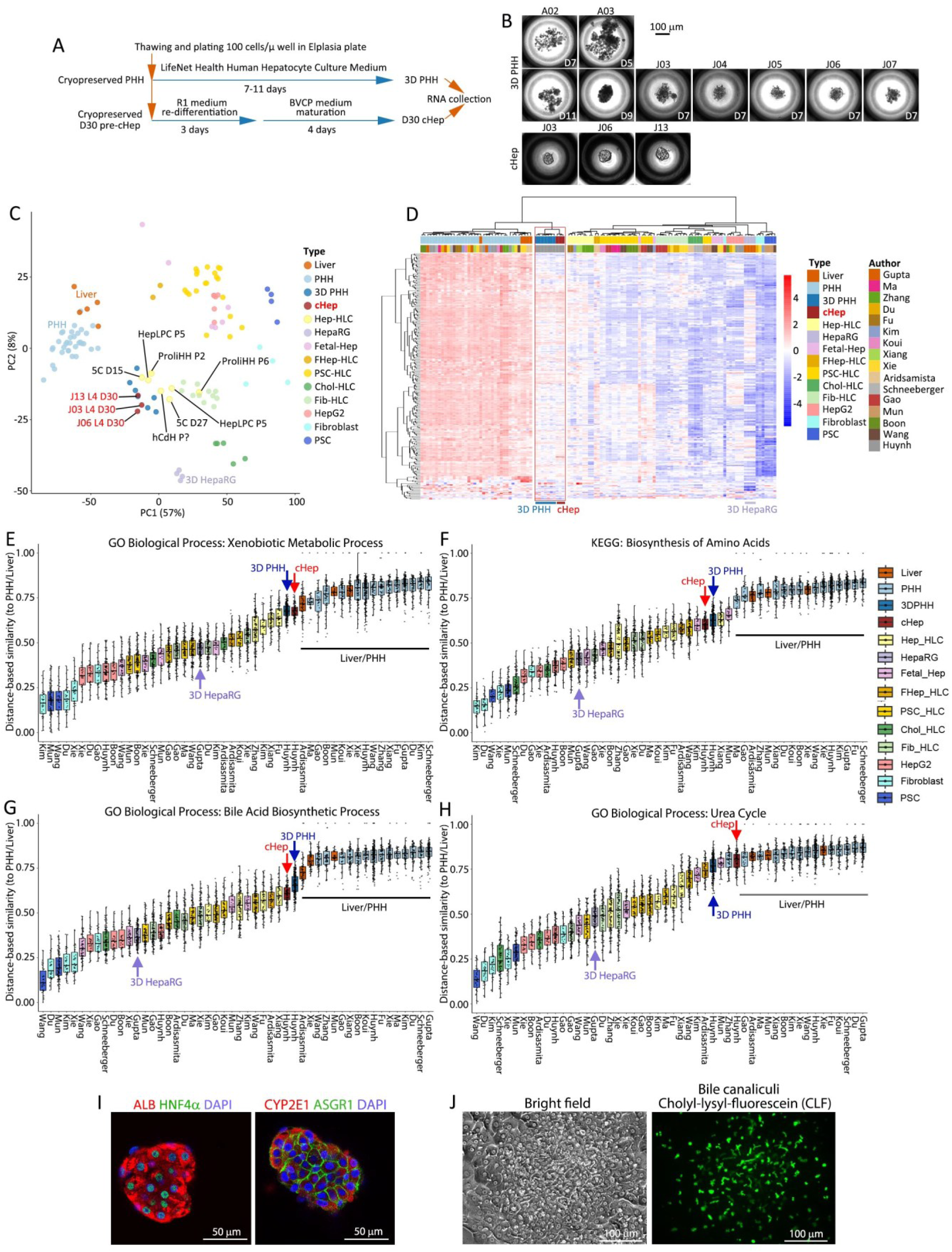
Characterization of cHep. **A.** Direct differentiation of cryopreserved pre-cHep using a protocol equivalent to 3D PHH spheroid formation. **B.** Representative images of 3D PHH and cHep. **C.** Principal component analysis (PCA) plot including cHep generated from day 30 (D30) J03, J06, and J13 L4 pre-cHep, 3D PHH, and published hepatocyte-like cells (HLC). **D.** Heatmap showing expression of 264 liver-enriched genes. **E–H.** Distance-based similarity scores based on gene expression associated with xenobiotic metabolism (E), amino acid biosynthesis (F), bile acid biosynthesis (G), and the urea cycle (H). **I.** Immunofluorescence staining for albumin, HNF4α, CYP2E1, and ASGR1. **J.** Cholyl-lysyl-fluorescein (CLF) staining.

### Genetically modified pre-cHep/cHep as a novel tool for disease modelling

The robust proliferation and preservation of differentiation capacity in pre-cHep enable the generation of genetically modified hepatocytes. As an example, we knocked out PNPLA3, whose rs738409 variant causes the I148M amino acid substitution and is strongly associated with metabolic dysfunction-associated steatotic liver disease (MASLD). Electroporation of Cas9/gRNA ribonucleoprotein (RNP) together with a plasmid carrying a puromycin resistant gene expression cassette, followed by transient puromycin selection (Figure 4A), achieved more than 89% of +1 bp insertion mutations in exon 3 across all tested pre-cHep lines (J03, J04, and J06) (Figures 4A and 4B). This process did not affect proliferation or differentiation capacity of pre-cHep (Figures 4C and 4D). When the edited cells were differentiated into cHep and cultured in the presence of free fatty acids, PNPLA3-deficient J03 and J04 cHep showed enhanced triglyceride accumulation, consistent with prior reports in iPSC-derived hepatic models and fetal hepatocyte organoids^32,33^, but not J06 cHep. These findings indicate that the effects of PNPLA3 loss are influenced by genetic background, and that pre-cHep/cHep derived from multiple donors may provide a powerful platform for studying MASLD susceptibility and developing new therapies.

**Figure 4.**
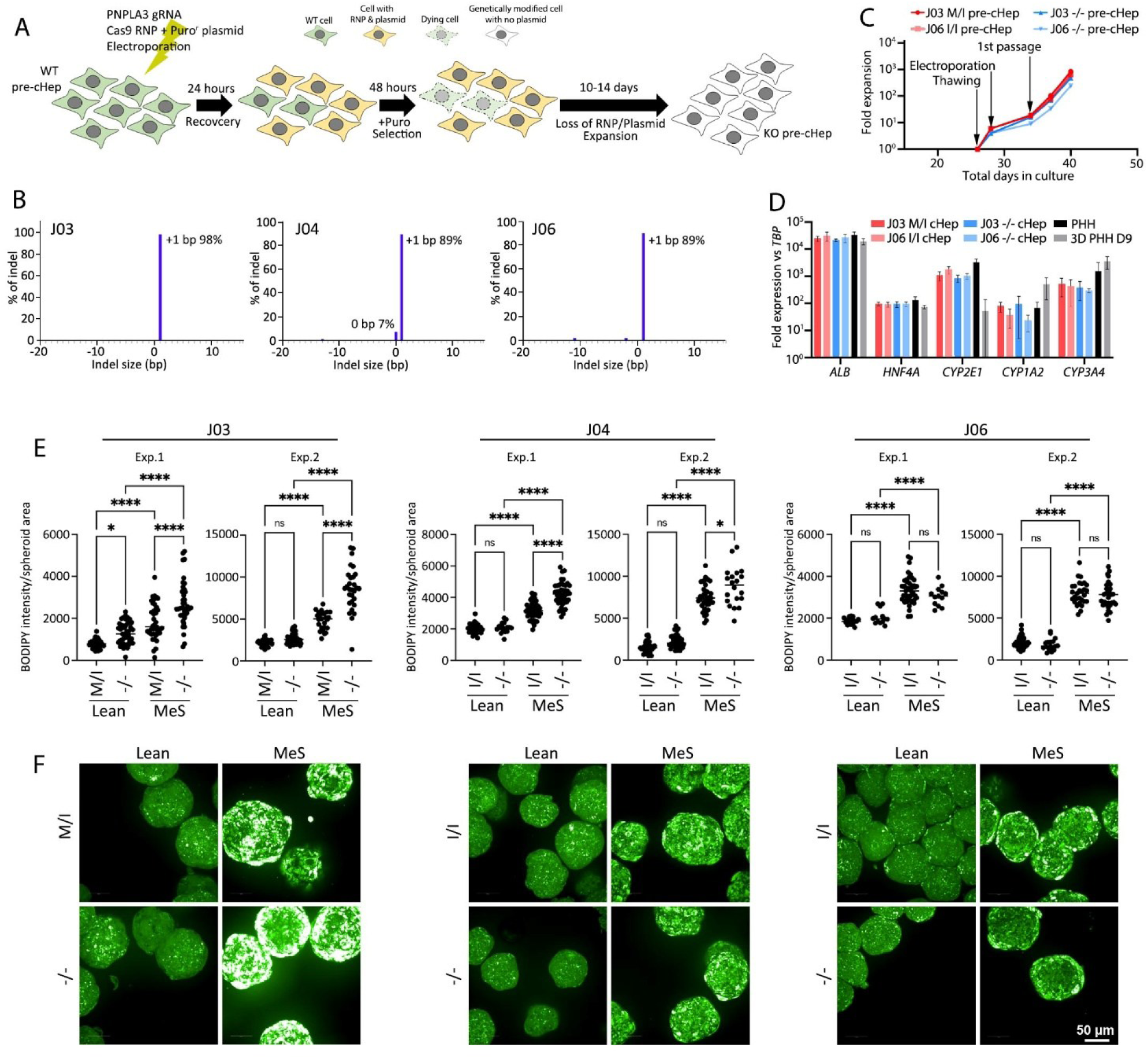
Generation of genetically modified pre-cHep/cHep. **A.** Strategy for generating a bulk PNPLA3⁻/⁻ pre-cHep population. **B.** Percentage of genetically modified cells estimated by ICE analysis. **C.** Proliferation of pre-cHep during genetic modification. **D.** RT-PCR analysis of hepatic gene expression in genetically modified cHep. **E.** Quantification of lipid accumulation detected by BODIPY staining in wild-type and PNPLA3⁻/⁻ cHep cultured under lean and metabolic syndrome (MeS) conditions for 10 days. **F.** Representative images of BODIPY staining.

In summary, L4 medium enabled the derivation of pre-cHep lines from juvenile PHH that expanded more than 10^6^-fold within 30 days while retaining liver repopulation capacity. cHep generated directly from cryopreserved pre-cHep as spheroids over 7 days showed global gene expression profiles similar to those of 3D PHHs and could provide a more affordable and reproducible alternative to PHHs for toxicology testing and disease modelling.

## Discussion

Hepatocytes are central to toxicology in drug development because they perform the majority of drug metabolism and generate metabolites that can be hepatotoxic. Cryopreserved primary human hepatocytes (PHHs) are widely used across industry, but batch-to-batch variability, limited donor supply and vial numbers, and high-cost complicate assay standardization. Our pre-cHep/cHep technology can expand PHH by more than 10^6^-fold within 30 days while retaining *in vivo* liver-repopulation and *in vitro* differentiation capacity, supplying a scalable source of high-quality human hepatocytes. Cryopreserved pre-cHep can be directly differentiated *in vitro* into 3D cHep spheroids in formats and timeframes compatible with existing industrial toxicology and metabolism workflows, simplifying adoption.

A current limitation is that this large-scale expansion retaining the functionality has been successful only with juvenile donor PHHs (donors aged ∼8 months to 10 years); nonetheless, 3 of 7 tested juvenile pre-cHep lines retained liver-repopulation capacity through day 30. The expansion magnitude is substantial: a single vial of juvenile PHHs could theoretically yield on the order of 10^6^ pre-cHep vials, which likely exceed current annual global usage of high-quality cryopreserved PHH. Notably, a single donor liver can yield up to 500-1,000 vials of cryopreserved PHH, and the majority of viable PHH that attach to culture plates subsequently proliferate in L4 medium.

For short-term ADME and acute toxicity assays (generally up to ∼48 hours), lower-cost, readily available cryopreserved non-platable PHHs remain adequate; however, for repeated-dose studies, slowly metabolized compounds, and chronic DILI assessment, longer exposures in 3D PHH models (spheroids, organoids, or long-lived co-cultures) are the preferred standard. pre-cHep/cHep supply addresses these needs by offering lower cost, larger supply, and greater lot consistency for longer-term and 3D assays. At the scale of producing ∼1000 vials of 1 million cells/vial from 1 million pre-cHep (i.e. 1,000-fold expansion), required total L4 medium costs just under £5000 (<£5/million pre-cHep/vial).

Overall, pre-cHep/cHep have the potential to replace many PHH spheroid-based assays and broaden access to human hepatocytes for toxicology, ADME, and disease modelling. Their scale and consistency can streamline assay optimization and standardization and even enable use of cells from the same donor across an entire multi-year drug development program, improving cost-effectiveness. Future priorities include expanding the donor range (improving L4), optimizing 2D differentiation and co-culture with non-parenchymal cells, and transitioning to GMP-grade processes—steps that would extend the technology’s utility into high-content screening, advanced disease models, and potentially cell therapy.

## Methods

### Generation and maintenance of pre-cHep lines

pre-cHep culture was carried out using small hepatocyte medium (SHM)^34^, comprised of DMEM/F12 (Sigma), 0.05% Bovine Serum Albumin (Gibco), 1:100 MEM Non-Essential Amino Acid Solution (Gibco), 10 ng/ml murine epidermal growth factor (EGF) (Peprotech), 1:100 insulin-transferrin-serine (ITS)-X (Life Technologies), 0.1 µM Dexamethasone (Sigma), 2.5 mM Nicotinamide (Sigma), 0.1 mM 2-phospho-L-ascorbic acid trisodium salt (Sigma), 2 mM L-glutamine (Invitrogen), 5 mM HEPES sodium salt (Gibco), and penicillin/streptomycin solution (Invitrogen) with following supplements. For L1, 5 µM Blebbistatin (ApexBio), 1.1 µM Vactosertib (MedChemExpress), 3 µM CHIR-99021 (Stem Cell Technologies), 12.5 ng/ml PG-001 (Peptigrowth), 10% fetal calf serum (FCS) were supplemented to SHM. For L2, 10 µM forskolin (Tocris) and 50 µM Dibutyl cyclic AMP sodium salt (db-cAMP) (Sigma) were supplemented to L1. For L4, 10 ng/ml human IL-6 (R&D Systems) was supplemented to L2. Nicotinamide was somewhat detrimental for a long-term expansion of some pre-cHep lines and not essential for all tested lines, thus omitted in some experiments. For the derivation of pre-cHep, cryopreserved primary human hepatocytes, listed in Supplemental Table S1, were seeded in L2 or L4 at 10^6^ cells per 10 cm plates, coated with 100 ug/ml rat tail collagen type I in PBS (Corning). The cells were cultured in a hypoxic condition with 3% O_2_, 5% CO_2_ at 37 °C with medium change every 2-3 days until the cells became confluent (6-10 days). The resulting pre-cHep was harvested with TrypLE and passaged in the same culture conditions. Typically, 5×10^4^ pre-cHep seeded in L2 and L4 in a well of a 6-well plate become 4-5×10^5^ in 4 days with one medium change on day 2, before reaching a total of 30 days in culture after which the proliferation speed in L2 slows down gradually. Cryopreservation of pre-cHep was carried out with CELLBANKER 2 (AMSBIO) at 1×10^6^/500 µl cells per cryovial, storing the vial directly in a −80 °C freezer.

### Enrichment of ASGR1+ pre-cHep

Isolation of ASGR1^+^ pre-cHep was carried out using poly-(N-p-vinylbenzyl-O-beta-D-galactopyranosyl-[1-4]-D-gluconamide) (PVLA)^35^ (Funakoshi). Briefly, 1 mg/ml PVLA was coated on a non-treated 6-well flat-bottom plate overnight at 37 °C. Subsequently, the wells were washed twice with PBS and then incubated with 1 ml/well 0.5% BSA in PBS at 37 °C for 30 minutes to avoid non-specific binding of ASGR1^−^ cells. After washing the wells twice with PBS, dissociated pre-cHep were seeded at 10^6^ cells/well and incubated at 37 °C. One hour later, unbound cells were removed by washing the wells twice with SHM containing 10% FCS. To collect PVLA-bound pre-cHep, the remaining cells were incubated in 1 ml of 0.02% EDTA in PBS for 5 minutes, collected in a centrifuge tube with 4 ml 10% FCS SHM, and then centrifuged at 300 g for 3 minutes. After removing the supernatant, the cells were resuspended in a culture medium for subsequent use.

### Differentiation of pre-cHep into cHep

In the experiments of Figure 1, pre-cHep seeded in Elplasia Microcavity plates (Corning) at 100 cells/microwell were cultured in L2 and cultured for 3 days. The resulting cHep spheroids were collected for various assays or maintained in the microwells in BVCP maturation/maintenance medium (BVCP) comprised of SHM supplemented with 5 µM Blebbistatin (ApexBio), 1.1 µM Vactosertib (MedChemExpress), 3 µM CHIR-99021 (Stem Cell Technologies), 1 µM PD0325901 (Sigma) under 3% O_2_, 5% CO_2_ at 37 °C. Later in more optimized differentiation protocol, pre-cHep (100 cells/microwell in Elplasia Microcavity plates or 1000 cells/well in BIOFLOA 96-well Cell Culture Spheroid pates) were cultured in re-differentiation medium R1 comprised of L2 medium supplemented with 1 ng/ml IL-6, 1 µM PD0325901, 1 µM Erlotinib, 5 µM DAPT, 2.5 µM LY294002, for 3 days under 20% O_2_, 5% CO_2_ at 37 °C. Subsequently, a 50% medium change was performed every 2 days using BVCP Maintenance/Maturation medium, consisting of SHM (containing 2.5 Mm nicotinamide) supplemented with 5 µM Blebbistatin, 1 µM Vactosertib, 3 µM CHIR99021, and 1 µM PD0325901, for at least two cycles, corresponding to a minimum 4-day maturation period before assays.

### ALBUMIN ELISA

Three days after differentiation of pre-cHep, the culture medium was changed to BVCP medium and then cultured for 1 day before being used for Albumin sandwich ELISA assay (Proteintech). The assay was conducted following the manufacturer’s instruction and the absorbance was measured using a Promega Glomax Explorer plate reader at 450 nm. The absorbance values were then compared to the standard curve generated to obtain the albumin concentration µg/day/million cells.

### CYP assay

CYP1A2, CYP2C9 and CYP3A4 activities were measured with P450-Glo™ CYP1A2, 2C9, and 3A4 assay kits (Promega) following the manufacturer’s instructions. Briefly, in a 96-well Elplasia plate, 100 pre-cHep per microwell were seeded for differentiation in the L2 medium. After 3 days, the spheroids were washed twice in 200 µl SHM. For the induction of CYP1A2, CYP2C9 or CYP3A4, the final concentration 100 µM Omeprazole (Sigma) or 1 mM Phenobarbital (Sigma) was supplemented in SHM. 48 hours later, the spheroids were washed twice in PBS with Mg^2+^ and Ca^2+^, and then incubated in 50 µl/well of CYPs substrate solution for 1 hour (CYP1A2, CYP3A4) or 4 hours (CYP2C9). After the incubation, 25 µl of CYPs substrate solution was transferred to an opaque white 96-well plate, and 25 µl of luciferin detection reagent was added to each well. After incubation for 20 minutes at room temperature in the dark, the luminescence was measured using a Promega Glomax Explorer plate reader. The relative luminescence unit (RLU)/ml/million cells was calculated after averaging the raw luminescence values from the technical duplicate wells and subtracting the background luminescence from the control wells.

### Differentiation capacity tests towards cholangiocytes

To test the differentiation capacity of pre-cHep to cholangiocytes, we have used two published protocols used for CLiP/CLiPLC^6,7^, with human biliary epithelial cells (hBECs) derived from cholangiocytes as a positive control^25^. In the method by Kim, 2×10^5^ hBECs or pre-cHep were re-suspended in 200 µl cholangiocytes differentiation medium (CDM), comprised of DMEM/F12 containing 10% FCS and 12.5 ng/ml PG-001 (Peptigrowth). The cell suspension was mixed with equal volume of CDM supplemented with collagen type 1 (BD Biosciences). The mixture is then transferred into a 6-well plate and incubate for 30 min at 37 °C until the gel was set. Thereafter, 2 ml of CDM was overlayed, and which was changed every 2 days. At day 7, cells were subjected to Rhodamine 123 transport assay harvested for RNA extraction. In the method by Katsuda, one day before seeding pre-cHep or hBECs, 5×10^4^ irradiated mouse fetal fibroblast (iMEF) were plated onto a collagen-coated 12-well plate. On the following day (Day 0), 5×10^5^ pre-cHep or hBECs were resuspended in SHM with 10 µM Y-27632, 0.5 µM A-83-01, 3 µM CHIR99021 (YAC) and 5% FCS and seeded on iMEF. Next day (Day1), medium was changed to mTeSR™1 with YAC (Stemcell Technologies) (mTeSR1+YAC). Medium was changed every 2 days. At day 7, the medium was replaced with mTeSR1+YAC supplemented with 2% Matrigel. Thereafter, medium was changed every 2 days. At day 14, cells were subjected to Rhodamine 123 transport assay or harvested for RNA extraction. Rhodamine 123 transport assay was carried out by incubating the cells with 100 µM of Rhodamine 123 (Sigma-Aldrich) for 5 minutes at 37 °C, washed 3 times with media and then kept in fresh media for 1 hour. After 1 hour, the cells were washed twice with media before imaging.

### Generation of liver humanized mice with TK-NOG-hIL6 mice

TK-NOG-hIL6 mice were housed at a room temperature of 24 °C with a 12 h light/dark cycle, with free access to irradiated (30 kGy) pellet diet (CLEA Rodent Diet CA-1; CLEA Japan, Tokyo, Japan) and tap water^24^. 7- to 8-week-old mice were given 0.1 to 0.8 mg/mL Val-ganciclovir (ValGCV; Sigma-Aldrich, St. Louis, MO, USA) in drinking water for 2 days to damage hepatocytes expressing the HSVtk transgene. Seven days after the administration of ValGCV, alanine aminotransferase (ALT) levels in plasma were measured using an automated clinical chemistry analyzer (Fuji Dri-Chem 7000; Fuji Photo Film, Tokyo, Japan). Liver-injured mice with ALT levels exceeding 400 U/L received pre-cHep transplantation. A total of 1.0 million cells pre-cHep in 50 µL of Williams’ medium E (Thermo Fisher Scientific) were intrasplenically injected using a 1/2-mL insulin syringe with a permanently attached needle (29 G × 0.5 inch; Terumo Corporation, Tokyo, Japan). The reconstitution of chimeric human liver was estimated by measuring plasma butyrylcholinesterase activity using an automated clinical chemistry analyzer^36^. The study protocol was reviewed and approved by the Animal Care Committee of Central Institute for Experimental Medicine and Life Science, and all experiments were conducted in strict accordance with the Guide for the Care and Use of Laboratory Animals (permit numbers: AIA260059).

### Immunohistochemistry of the liver

The chimeric livers from pre-cHep engrafted mice were fixed in 4% (v/v) phosphate-buffered formalin (Mildform 10NM; Wako Pure Chemical Industries), and 5-µm paraffin-embedded sections were prepared. For the immunohistochemical staining, some sections were autoclaved for 10 min in a target retrieval solution (0.1 M citrate buffer, pH 6.0; 1 mM EDTA, pH 9.0) and equilibrated at room temperature for 20 min. Primary antibodies included monoclonal mouse anti-human mitochondria antibody (monoclonal mouse anti-human mitochondria, clone 113-1, Merck Millipore, Burlington, MA), anti-HLA class I-A, B, C (HLA) (clone EMR8-5, Hokudo Co. Ltd., Sapporo, Japan), anti-human CYP2C9 (clone 2C8, LifeSpan Biosciences, Inc., Seattle WA. USA), and rabbit anti-human CYP3A4 (clone EPR6202, Abcam plc.). Mouse and rabbit Ig antibodies were visualized using amino acid polymer/peroxidase complex-labelled antibodies [Histofine Simple Stain MAX PO (MULTI); Nichirei Biosciences Inc.] and diaminobenzidine (DAB; Dojindo Laboratories, Kumamoto, Japan) substrate (0.2 mg/mL 3,3′-diaminobenzidine tetrahydrochloride in 0.05 M Tris-HCl, pH 7.6 and 0.005% H_2_O_2_). Sections were counterstained with hematoxylin. Images were captured using a digital slide scanner, NanoZoomer S60 (HAMAMATSU Photonics KK, Hamamatsu, Japan).

### Copy Number Variation (CNV) analysis

Genomic DNA was isolated from J06 pre-cHep at day 10 and day 60 of expansion, and CNV analysis was performed using the Infinium CytoSNP-850K BeadChip. The resulting data were analyzed with BlueFuse Multi software, following the method described by Steventon-Jones et al^37^.

### RNA-seq data processing and integration analysis

Raw paired-end RNA-seq FASTQ files from this study and publicly available datasets were processed using a unified analysis pipeline. Reads were aligned to the human reference genome GRCh38 primary assembly using STAR (v2.7.11b)^38^, and aligned BAM files were sorted and indexed using SAMtools (v1.20)^39^. Gene-level read counts were generated using featureCounts (v2.0.6)^40^. For samples sequenced across multiple lanes, reads were jointly aligned during STAR processing.

### Normalization and batch correction

Count matrices were imported into DESeq2 (v1.42.1) in R^41^. Variance stabilizing transformation (VST) was applied following DESeq2 normalization. Batch correction was performed using removeBatchEffect() from the limma package^42^.

### Principal component, Gene signature heatmaps

PCA was performed using the plotPCA() function in DESeq2 on variance-stabilized expression matrices. Liver gene sets, including the 264 liver-enriched genes, KEGG pathway-associated gene sets (biosynthesis of amino acids, retinol metabolism, fatty acid metabolism, and complement/coagulation cascades), and Gene Ontology biological process gene sets (urea cycle, xenobiotic metabolic process, and bile acid biosynthetic process), were used for expression analyses. Heatmaps were generated from variance-stabilized and batch-corrected expression matrices using pheatmap (v1.0.12). Hierarchical clustering was performed using the ward.D2 method. Dendrogram branches were reordered using the dendextend package for visualization without altering clustering relationships^43^.

### Distance-based similarity (DBS) analysis

Distance-based similarity analysis was performed following the approach described in HLCompR. Briefly, Euclidean distances were calculated from corrected expression matrices for predefined liver gene sets using primary human hepatocyte (PHH) samples as reference controls. Distances were transformed into similarity scores using the formula: *DBS=(D_max_-D_sample_)/D_max,_* where *D_sample_* represents the Euclidean distance of each sample to the PHH reference group, and *D_max_* represents the maximum observed distance within the dataset for a given gene set. Higher DBS scores indicate greater transcriptional similarity to PHHs.

### Genetic manipulation of pre-cHep

PNPLA3 knockout pre-cHep lines were generated using CRISPR/Cas9. For each electroporation, 125 pmol of gRNA targeting exon 3 (5’-GGGATAAGGCCACTGTAGAA-3’) was combined with 122 pmol of Cas9 in a sterile RNase-free PCR tube and incubated at room temperature for 15 minutes before being placed on ice. In the meantime, the cells were washed with PBS, and harvested with TrypLE at 37 °C for 5-10 minutes. 1×10^6^ cells were transferred to a sterile 1.5 ml tube and washed twice with Opti-MEM. Cells were gently resuspended in 100 µl of Opti-MEM containing the Cas9/gRNA complex and 10-20 µg of puromycin registrant gene expression plasmid, then transferred into an electroporation cuvette. Cell were electroporated using NEPA21 with the following settings: Poring pulse: voltage=200 V, pulse length=5 ms, pulse interval=50 ms, number of pulses=2, decay rate=10%, Polarity=+ and transfer pulse: voltage=20 V, pulse length=50 ms, pulse interval=50 ms, number of pulses=5, decay rate=40%, Polarity=+/-. Cells were left in a cuvette at room temperature for 45 minutes before being transferred into a single well of a 6-well plate. Cells were cultured in L4 medium for 24 hours before adding 5 μg/ml of puromycin. After subsequent 48 hours of puromycin selection, cells were expanded in L4 medium until becoming semi-confluent before passaging and genome collection for genotype check with ICE assay.

### Genotyping with ICE assay

321 bp genomic region flanking the PNPLA3 gRNA target site was amplified with primer PNPLA3F (5’-AAGTTTGTTGCCCTGCTCAC-3’) and PNPLA3R (5’-CCAGCTGTGGCTACTCTGTC - 3’), DreamTaq polymerase (Thermo Fisher) and 100 ng of genomic DNA using the following PCR program: 3 min at 95 °C (1x), followed by 20 sec at 95 °C, 30 sec at 58 °C, 30 sec at 72 °C (30x) and finishing with 2 min at 72 °C (1x) and incubation at 4 °C. Amplicons were column purified using the Zymoclean DNA clean and concentrate kit (Zymoclean). DNA concentration was measured using nanodrop. 50 ng of the amplicon was sent for Sanger Sequencing at Source Bioscience, using the forward amplification primer as sequencing primer. Chromatographs from edited and wild-type cells were uploaded on ICE website to analyse indel efficiency.

### In vitro steatosis model with cHep

pre-cHeps were plated at a density of 100-200 cells/microwell in an Elplasia plate to form spheroids in L2. After 3 days of differentiation, the medium was switched to lean or lipogenic BVCP. Lean BVCP contained 5.5 mM glucose and 172 pM insulin, with all other components in BVCP at the same concentrations. lipogenic BVCP contained 320 µM FFA (213 µM of oleic acid and 107 µM palmitic acid bound to BSA), 22 mM glucose and 1.7 µM insulin, with all other components in BVCP at the same concentrations. Media was changed on day 3, 5 and 7, and the spheroids were collected to a V-bottom plate precoated with 10 % FCS in DPBS (-) to prevent any spheroids adhesion to the plasticware on day 10. They were fixed with 4 % PFA at RT for 30 minutes and washed three times with DPBS (-). For lipid quantification, the spheroids were stained with 2.5 µM boron-dipyrromethene (BODIPY) and 10 µg/mL Hoescht 3343 in DPBS (-) overnight at 4 °C. The following day, they were washed 5 times and resuspended in clearing solution made from 2.5 M fructose (Sigma) and 60 % glycerol (Sigma, G9012). The spheroids were pipetted inside a circular imaging spacer placed on a 0.1 mm microscopy slide (SunJin Lab Co). Quantification was performed using Harmony software. Images were segmented based on nuclei staining. BODIPY intensity was quantified on max projections and normalised to the spheroid size.

## Acknowledgements

We thank PhoenixBio for the evaluation of pre-cHep liver repopulation capacity in the initial phase of the project, Mr. Kohei Nakazono and Ms. Noriko Suzuki for their assistance at CIEM. This work was supported by MRC-AMED Regenerative Medicine and Stem Cell Research Initiative (MR/V005537/1 to K.K. and 22bm0704056 to K.Y.) and the Cooperative Research Program (Joint Usage/Research Center program) of Institute for Life and Medical Sciences, Kyoto University (K.K. and K.Y.). JJ was supported by Martin Lee Doctoral Scholarship Programme at The University of Edinburgh. CTL and SKT were supported by the Research Impact Fund (Ref. No. 2410038 and 311003) and Collaborative Research Fund (Ref. No. 230094 and 4443560) of the Hong Kong Research Grants Council.

## Author Contributions

LMH. YH, JJ, IB, RG, SU, FK, VLG and KK designed and performed experiments. CTL contributed to the analyses of the RNA-seq data sets. TYM and SJF provided hBEC. SJF, KY, SKT, HS and KK obtained funding, supervised experiment design and data interpretation, and contributed to the manuscript preparation with LMH.

## Additional Information

Supplementary Information is available for this paper. Correspondence and requests for materials should be addressed to. A UK patent application covering pre-cHep culture conditions was filed in January 2025 (application number 2500890.5), followed by an international PCT filing in January 2026 (PG451002WO) to secure broad territorial protection.

---

**Extended Figure 1.**
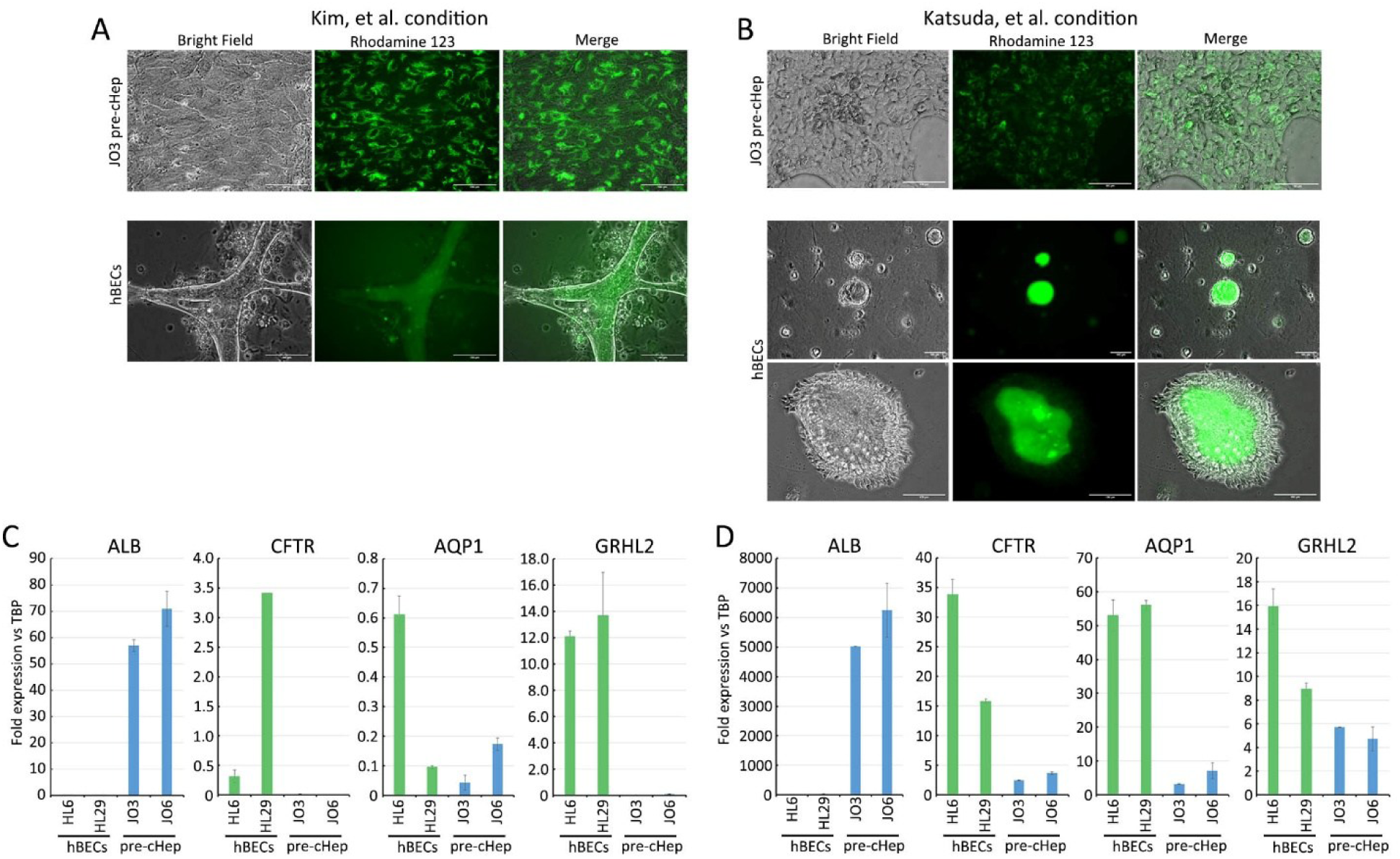
pre-cHep cultured under cholangiocyte differentiation conditions. **A.** Rhodamine 123 staining after 7 days of culture of pre-cHep under published CLiP/CLiPLC differentiation conditions toward cholangiocytes. Human biliary epithelial cells (hBECs) were used as a positive control. **B.** RT-PCR analysis of a hepatocyte marker ALBUMIN and cholangiocyte markers (CFTR, AQP1, and GRHL2) using samples from panel A. Data and error bars represent the mean ± SD from two technical replicates.

**Extended Figure 2.**
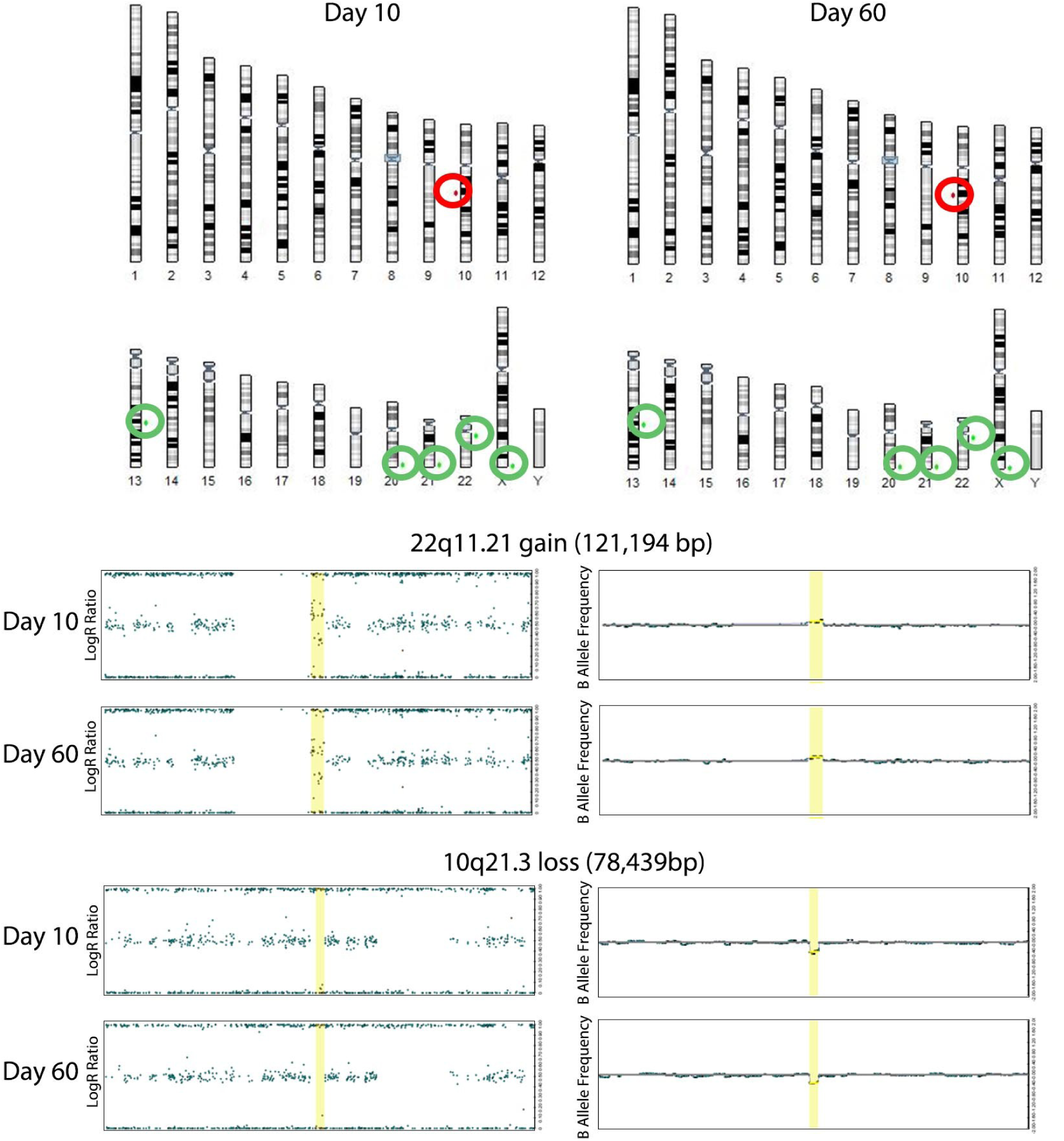
Copy number variation (CNV) in L4 pre-cHep at day 10 and day 60 of expansion. Gains and losses at specific loci relative to the reference baseline are mapped onto chromosomes and shown as red and green dots, respectively. No significant differences between the two time points were detected, indicating genetic stability during prolonged culture. Representative examples of a gained locus at 22q11.21 and a lost locus at 10q21.3 are shown.

**Extended Figure 3.**
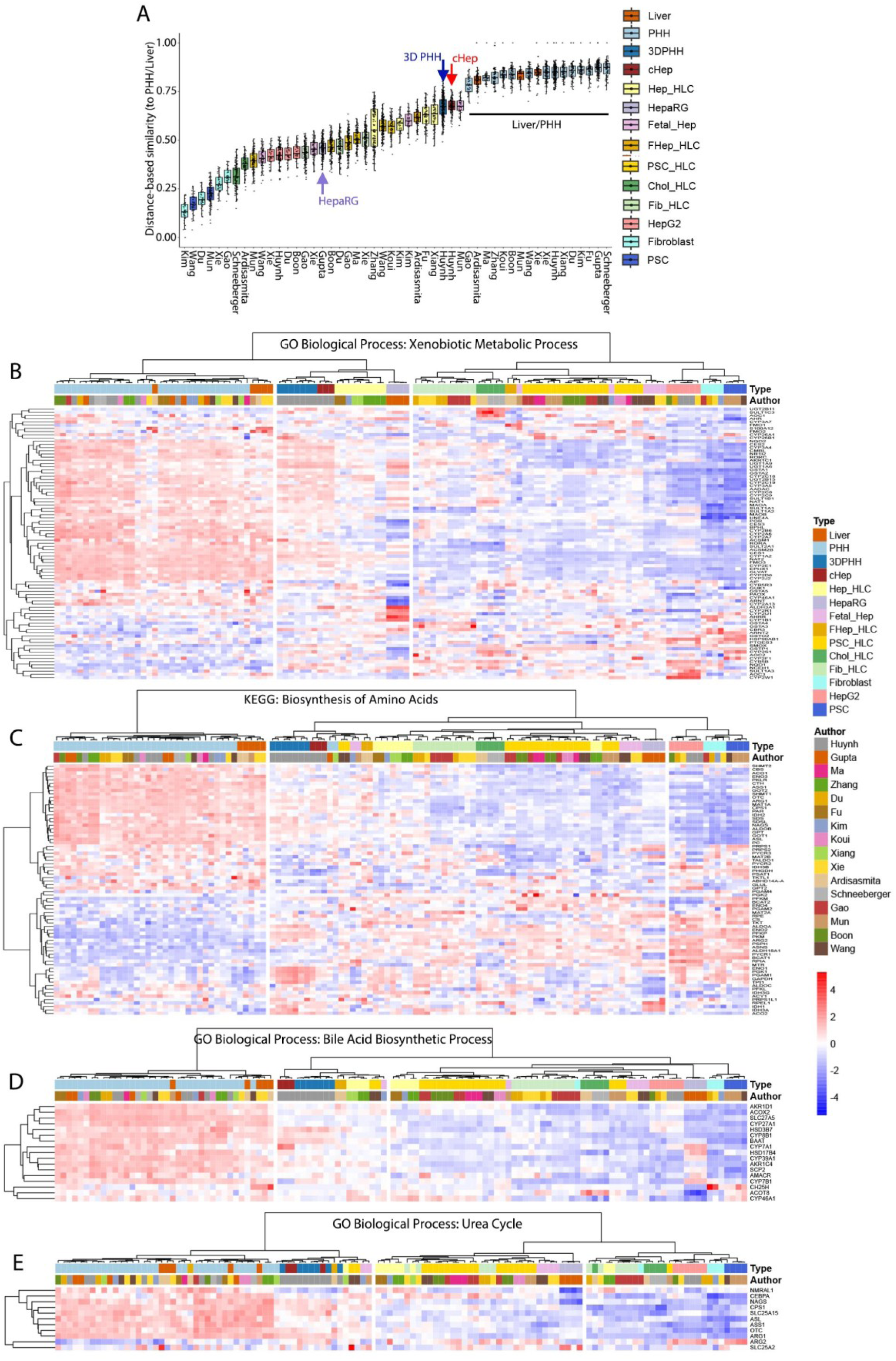
Gene expression analysis of HLC. Distance-based similarity score with 264 liver enriched genes (A), corresponding to Figure 3D. Heatmaps showing expression of genes involved in xenobiotic metabolism (B), amino acid biosynthesis (C), bile acid biosynthesis (D), and the urea cycle (E), corresponding to Figure 3E.

**Extended Figure 4.**
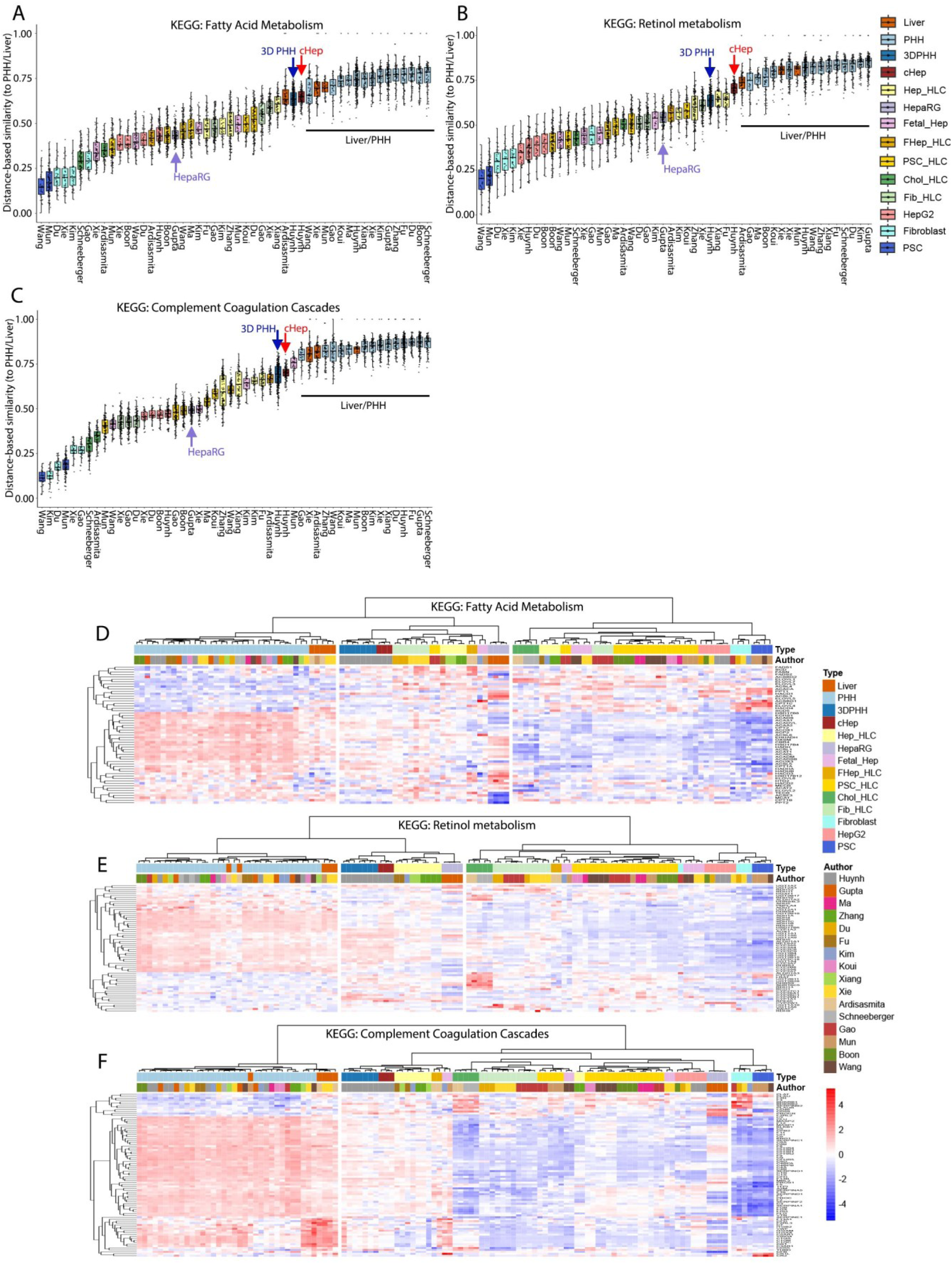
Gene expression analysis of HLC. Distance-based similarity score and gene expression heatmap with KEGG gene sets: Fatty acid metabolism (A, D), Retinol metabolism (B, E) and Complement coagulation cascade (C, F).

